# Treatment-Specific Composition of Gut Microbiota Is Associated with Disease Remission in a Pediatric Crohn’s Disease Cohort

**DOI:** 10.1101/412890

**Authors:** Daniel Sprockett, Natalie Fischer, Rotem Sigall Boneh, Dan Turner, Jarek Kierkus, Malgorzata Sladek, Johanna C. Escher, Eytan Wine, Baruch Yerushalmi, Jorge Amil Dias, Ron Shaoul, Michal Kori, Scott B. Snapper, Susan Holmes, Athos Bousvaros, Arie Levine, David A. Relman

**Affiliations:** Department of Microbiology & Immunology, Stanford University School of Medicine, Stanford, CA 94305, USA; Division of Infectious Diseases & Geographic Medicine, Department of Medicine, Stanford University School of Medicine, Stanford, CA 94305, USA; Pediatric Gastroenterology and Nutrition Unit, Wolfson Medical Center, Holon, Israel; The Juliet Keidan Institute of Pediatric Gastroenterology & Nutrition, Shaare Zedek Medical Center, The Hebrew University of Jerusalem, Jerusalem, Israel; Department of Gastroenterology, Hepatology, Feeding Disorders and Pediatrics, The Children’s Memorial Health Institute, Warsaw, Poland; Department of Pediatrics, Gastroenterology and Nutrition, Jagiellonian University Medical College, Cracow, Poland; Department of Pediatric Gastroenterology, Erasmus MC-Sophia Children’s Hospital, Rotterdam, The Netherlands; Division of Pediatric Gastroenterology and Nutrition, Department of Pediatrics, University of Alberta, Edmonton, Canada; Pediatric Gastroenterology Unit, Soroka University Medical Center, and Faculty of Health Sciences, Ben-Gurion University of the Negev, Beer Sheva, Israel; Department of Pediatrics, Hospital de Sao Joao, Porto, Portugal; Pediatric Gastroenterology Unit, Ruth Children’s Hospital, Rambam Medical Center, Haifa, Israel; Pediatric Day Care Unit, Kaplan Medical Center, Rehovot, Israel; Division of Gastroenterology, Hepatology, and Nutrition, Boston Children’s Hospital, Boston, MA 02115, USA; Division of Gastroenterology, Brigham and Women’s Hospital, and Harvard Medical School, Boston, MA, 02115, USA; Department of Statistics, Stanford University, Stanford, CA 94305, USA; Sackler School of Medicine, Tel Aviv University, Tel Aviv, Israel; Infectious Diseases Section, Veterans Affairs Palo Alto Health Care System, Palo Alto, CA 94304, USA

**Keywords:** pediatric Crohn’s disease, microbiota, antibiotics, disease remission, Random Forest model

## Abstract

**Background:** The beneficial effects of antibiotics in Crohn’s disease (CD) depend in part on the gut microbiota but are inadequately understood. We investigated the impact of metronidazole (MET) and metronidazole plus azithromycin (MET+AZ) on the microbiota in pediatric CD, and the use of microbiota features as classifiers or predictors of disease remission.

**Methods:** 16S rRNA-based microbiota profiling was performed on stool samples from 67 patients in a multinational, randomized, controlled, longitudinal, 12-week trial of MET vs. MET+AZ in children with mild to moderate CD. Profiles were analyzed together with disease activity, and then used to construct Random Forest models to classify remission or predict treatment response.

**Results:** Both MET and MET+AZ significantly decreased diversity of the microbiota and caused large treatment-specific shifts in microbiota structure at week 4. Disease remission was associated with a treatment-specific microbiota configuration. Random Forest models constructed from microbiota profiles pre- and during antibiotic treatment with metronidazole accurately classified disease remission in this treatment group (AUC of 0.879, 95% CI 0.683, 0.9877; sensitivity 0.7778; specificity 1.000, *P* < 0.001). A Random Forest model trained on preantibiotic microbiota profiles predicted disease remission at week 4 with modest accuracy (AUC of 0.8, *P* = 0.24).

**Conclusions:** MET and MET+AZ antibiotic regimens in pediatric CD lead to distinct gut microbiota structures at remission. It may be possible to classify and predict remission based in part on microbiota profiles, but larger cohorts will be needed to realize this goal.

**Summary:** We investigated the impact of metronidazole and metronidazole plus azithromycin on the gut microbiota in pediatric Crohn’s disease. Disease remission was associated with a treatment-specific microbiota configuration, and could be predicted based on pre-antibiotic microbiota profiles.

## Introduction

The global incidence of Crohn’s disease (CD) has steadily risen over the past few decades, especially among pediatric patients.^1^ Early diagnosis as well as timely and effective personalized treatment are especially crucial in this vulnerable patient population, as the disease in childhood may be more extensive and may follow a more severe course, with children experiencing growth failure or delayed puberty and life-long consequences. ^2,3^ Large efforts are being made to identify novel non-invasive biomarkers in CD patients, not only to monitor disease severity but also to predict therapeutic response.^4,5^

Genome-wide association studies have identified a variety of risk loci within genes important for the maintenance of homeostasis with our commensal gut microbiota.^6^ In this regard, several studies in pediatric CD patients have described a state of microbiome ‘dysbiosis’, with varying claims about disease-promoting or -ameliorating bacterial taxa^7–9^ and about differences in bacterial diversity.^10^ Antibiotics in general have a large effect on the microbiome, but the magnitude of the effect differs among individuals, as do taxa-specific responses to the same antibiotic treatment in different individuals.^11^ Similarly, several trials in adult CD have found, in general, a benefit from antibiotic treatment, but with heterogeneous results for the use of metronidazole, quinolones, and rifaximin.^12^

Metronidazole (MET) is one of the most commonly prescribed antibiotics in the treatment of pediatric CD, but recent studies reported superior outcomes for the combination of MET plus Azithromycin (AZ), as compared to MET alone.^13,14^ AZ is especially promising in the treatment of IBD, as it penetrates multiple intestinal compartments, including the intestinal lumen, biofilms in the mucus layer, as well as macrophages.^15,16^ The use of AZ might improve treatment of various pathogens implicated in CD, such as adherent and invasive *E. coli* (AIEC) strains.^17–19^

The goal of this study was to evaluate the intestinal microbiota as a source of predictors of disease state and of treatment response in pediatric CD patients, undergoing either single (MET) or combination (MET+AZ) antibiotic therapy. We took advantage of a randomized controlled trial of those two antibiotic regimens in pediatric CD and examined the treatment-specific impact on microbiome diversity and composition; we then employed machine-learning techniques to identify microbiota composition-based signatures associated with antibiotic treatment that might enable monitoring of disease and guide clinical decision-making.

## MATERIALS and METHODS

### Study Design

Seventy-four CD patients (ages 5-18 years) were previously enrolled at 11 pediatric gastroenterology clinical sites in an investigator-blinded randomized controlled trial comparing the efficacy of MET+AZ versus MET therapy for the treatment of children with mild to moderate active CD (10 ≤ Pediatric Crohn’s Disease Activity Index (PCDAI) ≤ 40) (National Institutes of Health NCT01596894). Sixty-seven of those 74 patients from 9 sites provided stool samples for the microbiome study reported here. A complete description of the study design, laboratory and analysis methods, and primary outcomes was published previously.^13^ Additional inclusion criteria, besides age and disease activity, included at least one elevated inflammatory marker above normal values (C-reactive protein, CRP; erythrocyte sedimentation rate, ESR; or calprotectin) and disease duration since diagnosis < 3 years. Exclusion criteria included presence of a stool pathogen (based on bacterial culture, parasite study, or *Clostridium difficile* toxin assay), involvement of the proximal ileum or jejunum (L4b as per Paris classification), unclassified IBD, fibrostenotic disease (defined as strictures with prestenotic dilatation), internal or perianal fistulizing disease, prominent extraintestinal manifestations (e.g., arthritis, uveitis and sclerosing cholangitis), known allergy to either metronidazole or azithromycin, prolonged QT_c_ at baseline, or steroid use during the 7 days prior to enrollment.

The treatment protocol was adapted from a report by Levine and Turner.^14^ Patients were enrolled in one of the two treatment arms with 1:1 randomization, although 11 subjects with lack of response to MET therapy received open-label AZ between weeks 4 and 8. These MET/MET+AZ subjects were treated as a separate group for analyses involving time points after week 4. Furthermore, 4 patients in the MET group, 5 patients in the MET+AZ group and 1 patient in the MET/MET+AZ group received steroids or biologics after week 4 at their physician’s direction due to inadequate response. Both MET and MET+AZ groups discontinued antibiotics after week 8, while the MET/MET+AZ group maintained their regimen to complete a total of 8 weeks. 67 patients participated in the microbiome study, where stool samples for microbiota analysis were collected at weeks 0, 4, 8 and 12. Disease activity was determined using the PCDAI and clinical remission was defined as PCDAI < 10.

### Study procedures for assessment of inflammatory markers

A blood sample for measurement of CRP, ESR, complete blood count and serum albumin was collected at each visit, as well as a fecal sample for measurement of calprotectin at weeks 0 and 8. Stool extracts were prepared using the BÜHLMANN Smart-Prep kit (Buhlmann Laboratories AG, Switzerland). Fecal calprotectin was measured using the fCAL ELISA kit (Buhlmann Laboratories AG, Switzerland) with a normal range <100μg/g. Preliminary results indicated very high levels of faecal calprotectin (>1800μg/g) in many patients; therefore, all samples were diluted 1:10 according to manufacturer’s instructions.

### Microbiota Analysis

Stool samples were collected and frozen on site at −20°C and then shipped to a central laboratory and stored at −80°C until further processing. DNA was extracted from ~200 mg stool using the QIAGEN DNeasy PowerSoil HTP 96 Kit (Qiagen, Cat #12955-4) following the manufacturer’s instructions, including a 2 × 10 minute bead-beating step using the Retsch 96 Well Plate Shaker at speed 20. The V4 region of the 16S rRNA gene was amplified using forward primer 515F (5’-GTGCCAGCAGCCGCGGTAA-3’) with an error-correcting barcode and 806R (5’-GGACTACCAGGGTATCTAAT-3’).^20^ PCR products were then purified, pooled in equimolar concentrations, and sequenced on three 2 × 300 Illumina MiSeq paired-end runs using reagent kit v3.

Raw reads were demultiplexed using the QIIME command `split_libraries_fastq.py` (QIIME version 1.9.1) and then quality trimmed using the DADA2 pipeline (dada2 version 1.1.1) in R (R version 3.2.4).^21^ Briefly, forward reads were truncated to 220 bp, and reverse reads to 150 bp, and quality-filtered with settings `maxN=0, maxEE=2, truncQ=2`. Following quality-trimming, amplicon sequence variants (ASVs) were inferred using the DADA2 pipeline. Taxonomy was assigned to each ASV using the RDP classifier and the SILVA 16S rRNA database (SILVA nr version 132), and a phylogenetic tree was built from the ASVs using the `phangorn` R package (`phangorn` version 2.4.0). The ASV table, patient sample data, taxonomy assignments, phylogenetic tree, and ASV sequences were then bundled into a single `phyloseq` data object for further plotting and statistical analysis (`phyloseq` version 1.24.2).^22^ Faith’s Phylogenetic Diversity was calculated using the R package `picante` (picante version 1.7), and distance calculations and ordination plots were built using the `phyloseq` package. Hierarchical clustering of ASVs was performed using the hclust function in the `stats` R package (stats version 3.5.1). Differential abundance tests were carried out using the `DESeq2` R package (DESeq2 version 1.20.0). Analysis of variance in the dataset was assessed with PERMANOVA (`vegan` version 2.5-4) with 1,000 permutations on the Unifrac distance matrix. Bonferroni correction was applied for multiple comparisons (`p.adjust` function in the `stats` package version 3.5.1).

### Random Forest Modeling

Machine learning analyses were performed using the `caret` and `randomForest` R packages (`caret` version 6.0.80, `randomForest` version 4.6.14). Prior to model building, the ASV abundance was transformed using a variance stabilizing transformation to normalize for sequencing depth in the `DESeq2` R package, as well as centered to reduce the influence of inter-individual variation. Models were constructed with samples from subjects in the MET group only. Model 1 was constructed on microbiota data from weeks 4 and 8, and Model 2 was constructed on microbiota data from weeks 0, 4 and 8, with the outcome variable being disease state (i.e., remission or non-remission) at the time of sample collection. Each model was trained on samples from a random subset of 70% of subjects, with generation of 1000 trees and leave-one-out cross-validation. To assess their accuracy, models were used to predict remission or non-remission on samples from the remaining 30% of subjects. Both sets of classification results were then evaluated by calculating their area under the curve (AUC), as derived from a receiver operating characteristic (ROC) curve analysis using the R package `ROCR` (`ROCR` version 1.0.7). Each model also calculated importance scores for ASVs based on the increase in prediction error when the ASV in question was left out of the training set via random permutation. Because the numbers of remission and non-remission samples in the MET+AZ group were too uneven to be able to construct a useful model, we focused on the MET group only.

A Random Forest model was constructed on all pre-antibiotic (week 0) microbiota data, as well as on clinical data, including sex, age, disease duration, tissue involvement (Paris classification^23^), baseline immunomodulators, and baseline CRP and PCDAI to predict disease state (remission or non-remission) at week 4 in both treatment groups combined. In order to increase the number of shared features found across samples and decrease the noise among closely-related bacterial taxa, ASVs were agglomerated based on their cophenetic distance in the phylogenetic tree at the `h = 0.3` level. These numbered ASV clusters were also labeled by the number of ASVs in the cluster, as well as the genus that was assigned to the majority of the ASVs within the cluster. If >50% of the ASVs within a cluster were not assigned to a single genus, the same majority rule was used iteratively at higher taxonomic levels. Models were constructed and tested as described above. Abundances were normalized using variance stabilizing transformation (`vst` function in the `DESeq2` package version 1.20.0).

Amplicon sequencing data are available at SRA (SRP160897). A detailed description of this analysis, along with all analysis code and raw data used to go from sequences to final figures, is available at the Stanford Digital Repository (https://purl.stanford.edu/mp935wb0227) and as Supplementary Data (see text, Supplementary Data Content 1).

## ETHICAL CONSIDERATIONS

All patients provided written informed consent, or assent when required, prior to participation. The study was conducted according to the principles of the Declaration of Helsinki.

## RESULTS

### Clinical Outcomes

A total of 67 patients from 9 sites provided stool samples for microbiome analysis and are included in the present study (Table 1). Of this group, 36 patients were randomly assigned to receive MET, and 31 to receive MET+AZ (Fig. 1A). Figure 1B shows the temporal dynamics of each patient’s PCDAI, measured at the time of stool sample collection (Fig. 1B). By week 4, 42% (15/36) of MET patients and 65% (20/31) of MET+AZ patients had achieved remission (Fig. 1B). Between weeks 4 and 8, 11 of the MET patients failing remission had azithromycin added to their treatment regimen (MET/MET+AZ). At week 8, 60% (15/25) patients in the remaining MET group, 65% (20/31) in the MET+AZ group, and 45% (5/11) in the new MET/MET+AZ group had achieved remission. At week 12, 72% (18/25) of the remaining MET group were in remission, while 71% (22/31) of the MET+AZ group and 100% (11/11) of the MET/MET+AZ group were in remission. Differences in remission rates and inflammatory markers between the data presented here (per protocol analysis) and the original clinical paper^13^ (intention-to-treat analysis) are due to the transfer of treatment failures out of the MET group to the MET/MET+AZ group, thus reducing the number of inflamed non responders in the MET group.

**FIGURE 1.**
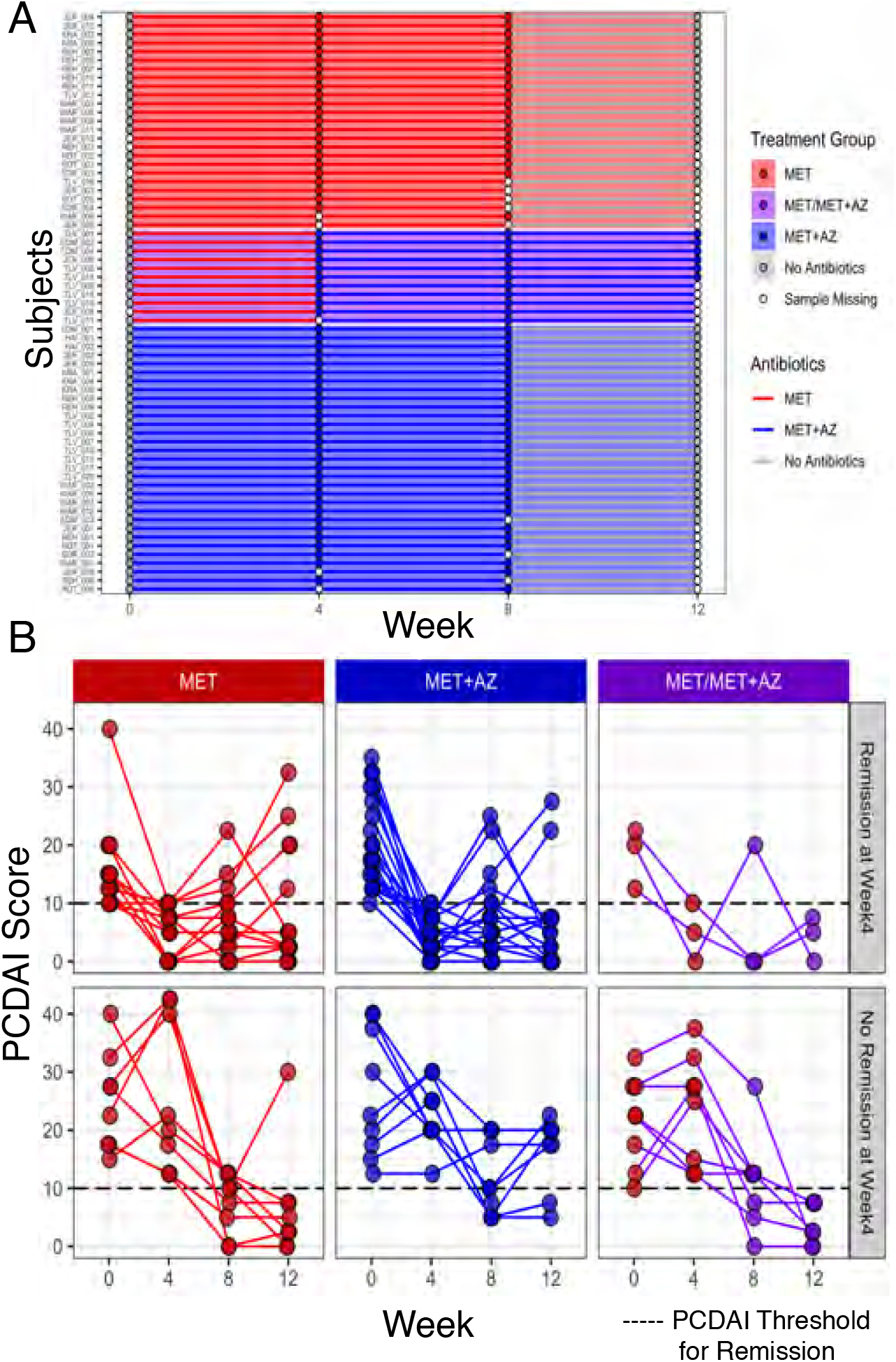
Study design and clinical outcomes. **A,** At week 0, 67 pediatric Crohn’s Disease patients were randomly assigned to one of two treatment groups: MET subjects received 20 mg/kg metronidazole twice daily (maximum of 1000 mg/day) for 8 weeks, while MET+AZ subjects received metronidazole plus 7.5 mg/kg azithromycin (maximum of 500 mg/day) once a day for 5 consecutive days, followed by a 2-day drug holiday, each week for the first 4 weeks and then stepped down to 3 consecutive days of the same dose with a 4-day drug holiday, per week over the subsequent 4 weeks. Metronidazole (MET, red, n = 36) or metronidazole and azithromycin (MET+AZ, blue, n = 31). Patients not in remission between weeks 4 and 8 could be offered open-label azithromycin based on physician assessment (MET/MET+AZ, purple, n = 11), and are displayed as a distinct patient cohort from weeks 4 to 12. Stool samples were collected at weeks 0, 4, 8, and 12. Stool samples are represented by a circle, while antibiotic treatment is represented by a line connecting the circles. **B,** Pediatric Crohn’s Disease Activity Index (PCDAI) for subjects at weeks 0, 4, 8, and 12. Treatment groups (MET, red; MET+AZ, blue; MET/MET+AZ, purple) are in columns, while the rows are groups of subjects that were either in remission (top) or not in remission (bottom) at week 4.

**Table 1.** Study subject features at baseline.

|  | **MET Group** | **MET+AZ Group** |
| --- | --- | --- |
| **Subjects** | 36 | 31 |
| **Samples** | 120 | 112 |
| **Female/Male** | 20/16 | 6/25 |
| **Mean Age (years) ±SD** | 13.5 ± 3.1 | 14.2 ± 3.1 |
| **Mean Disease Duration (years) ± SD** | 0.7 ± 1 | 1.1. ± 1.1 |
| **Baseline PCDAI ± SD** | 19.6 ± 8.1 | 22 ± 9 |
| **Disease Location (Paris Classification)** |  |  |
| **L1** | 13 | 6 |
| **L2** | 1 | 2 |
| **L3** | 22 | 23 |
| **L4a** | 18 | 16 |
| **Treatment at Baseline** |  |  |
| **Immunomodulators** | 14 | 19 |
| **Aminosalicylates** | 6 | 11 |
| **Antibiotics within the previous 12 months** | 2 | 2 |

67 pediatric Crohn's Disease patients were randomly assigned to treatment with either metronidazole (MET) or metronidazole plus azithromycin (MET+AZ). Gender (Chi-square test, $P < 0.01$ ) and number of subjects with L1 involvement (Chi-square test, $P < 0.01$ ) differed significantly between the two groups. *PCDAI* pediatric Crohn's Disease activity index. *SD* standard deviation.

Of note, our analysis included samples from 2 patients in the MET group and 2 patients in the MET+AZ group who did not respond to antibiotic treatment by week 4 and either received open-label steroids at week 4 or biologics at week 8. In addition, 6 patients in the MET group, 7 patients in the MET+AZ group and 3 patients in the MET/MET+AZ group received additional immunomodulators at some point during the study. Accordingly, not all clinical improvement in these patients could necessarily be attributed to the antibiotic regimen alone. Therefore, we examined the effect of additional drugs on achievement of remission. There was no difference between patients that started at baseline on either aminosalicylates (Chi-square test, *P* = 0.43) or immunomodulators (Chi-square, *P* = 0.10). Patients who received additional immunomodulators during the study were less likely to achieve remission (Chi-square test, *P* < 0.01). There was no difference between the numbers of patients who achieved remission after addition of steroids or biologics during the study and the numbers of patients achieving remission without additional drugs (Chi-square test: steroids *P* = 0.55, biologics *P* = 0.35).

Calprotectin, CRP and ESR are prominent markers of inflammation, currently measured in the clinic and interpreted by health care providers to determine disease status. CRP values decreased significantly in all three groups from weeks 0 to 4 of treatment (Wilcoxon Test, MET *P* = 0.0046, MET+AZ *P* = 3.2e-05, MET/MET+AZ *P* = 0.023), with no further significant changes from weeks 4 to 8 (see Supplementary Fig. 1A, as Supplementary Data Content 2). Fecal calprotectin measured was generally high (>1800 ug/g) at weeks 0 and 8, and decreased significantly in the MET+AZ (Wilcoxon test, *P* = 0.0093) and the MET/MET+AZ groups (Wilcoxon test, *P* = 0.0074), but not in the MET group (see Supplementary Fig. 1B, as Supplementary Data Content 2). We did not observe any significant changes in ESR levels following antibiotic treatment in any of the three groups (Wilcoxon Test, *P* <= 0.05), indicating that ESR might not be a good readout of disease status in this patient cohort (see Supplementary Fig. 1C, as Supplementary Data Content 2).

### Microbiota Response to Single and Combination Antibiotic Therapies

Antibiotics can strongly perturb the distal gut microbiota in an individual-specific manner;^24^ individualized responses might explain some of the observed heterogeneity in disease progression and dynamics. First, we characterized the microbiome of all patients at baseline. We calculated the mean Faith’s phylogenetic diversity and observed no differences between the three treatment groups at baseline (Supplementary Fig. 2A). Next, we calculated the mean unweighted Unifrac pairwise distance among all samples and did not observe significant differences in microbiome composition between the treatment groups at baseline (PERMANOVA, corrected *P*-value = 1) (Supplementary Fig. 2B). Since gender and number of subjects with L1 Paris Classification were unequally distributed between the MET and MET+AZ groups at baseline, we also tested for potential influence of these factors on baseline microbiome composition and detected no difference between male and female patients (PERMANOVA, corrected *P*-value = 1) (Supplementary Fig. 2C) or by L1 Paris Classification status (PERMANOVA, corrected *P*-value = 0.25) (Supplementary Fig. 2D). Patients with baseline use of immunomodulators or aminosalicylates had a stable disease state at the start of the study and were not excluded, and continued their medication throughout.

Aminosalicylates had a significant impact on baseline composition (PERMANOVA, corrected *P*-value < 0.05, Supplementary Fig. 2E), but immunomodulators had no impact on baseline microbiome composition (PERMANOVA, corrected *P*-value = 0.13, Supplementary Fig. 2F). Since the distribution of patients receiving 5ASA was similar in each of the treatment groups, we did not exclude those patients from the analysis.

We observed a profound shift in gut microbiota diversity and structure following antibiotic administration in most subjects. In the MET and MET+AZ treatment groups, alpha diversity decreased significantly (Wilcoxon test, *P* ≤ 0.01) (Fig. 2A). Diversity remained low in both groups at week 8, and rebounded towards its pre-antibiotic level by week 12, after discontinuation of antibiotics. Yet, only the MET group achieved a level of diversity that was comparable to its baseline state (MET week 0 vs. 12, Wilcoxon test, *P* = 0.16), while the MET+AZ group’s mean diversity remained significantly lower (MET+AZ week 0 vs. 12, Wilcoxon test, *P* ≤ 0.01).

**FIGURE 2.**
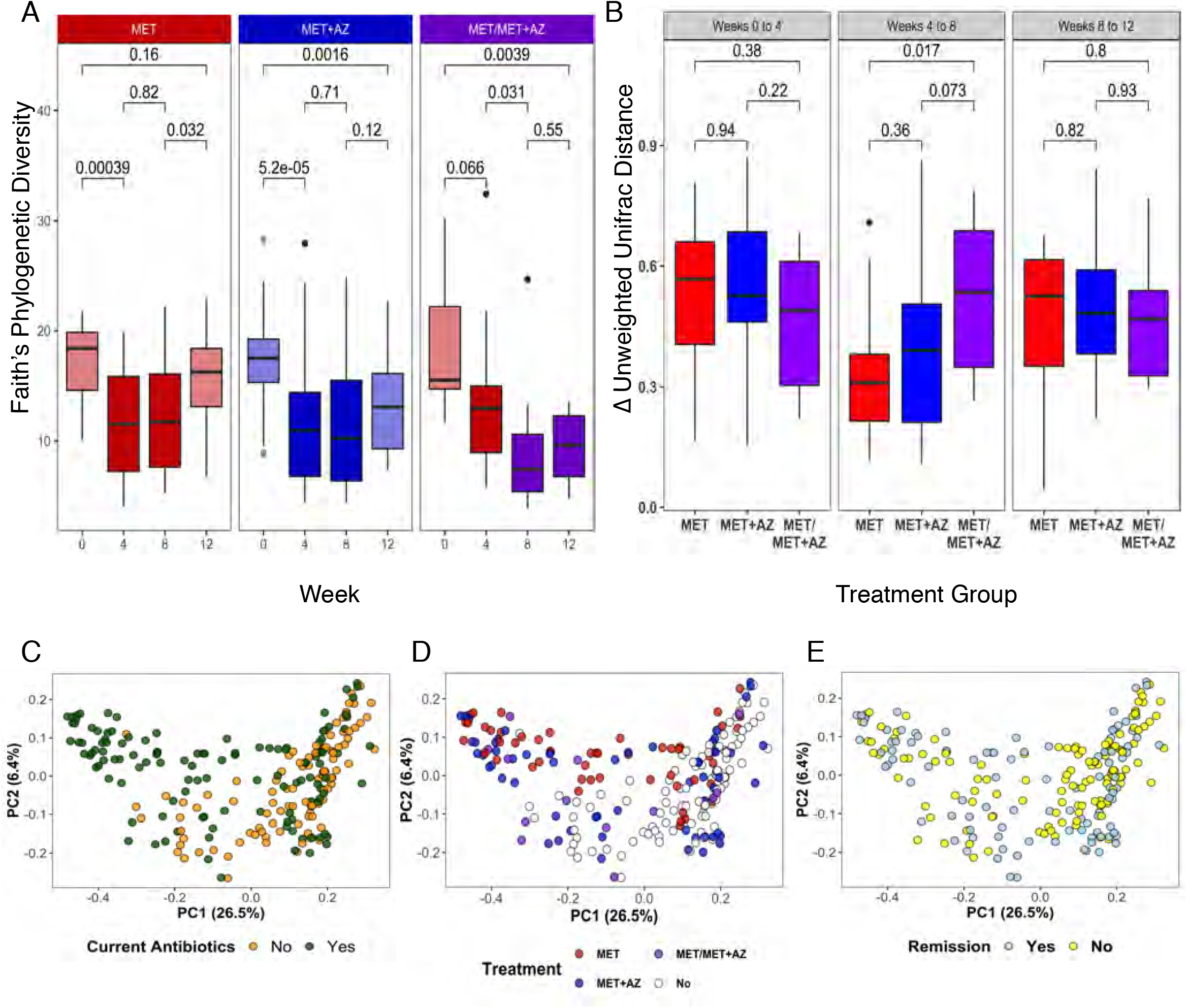
Microbiota response to antibiotic therapy. A, Alpha diversity measured by Faith’s Phylogenetic Diversity, segregated by treatment group (MET, red; MET+AZ, blue; MET/MET+AZ, purple) and week. Treatment group colors are darker during antibiotic treatment. *P*-values are displayed for comparisons between treatments (Wilcoxon Test). B, Boxplot of the median intra-subject unweighted UniFrac pairwise distance among samples collected during consecutive time points. C-E, Principal coordinates analysis (PCoA) on the unweighted UniFrac pairwise distances among all samples and all subjects. Samples are colored by C, current antibiotic use, yes/no (PERMANOVA, *P* < 0.01); D, current antibiotic use and treatment group (PERMANOVA, *P* = 0.1) and remission status (PERMANOVA, *P* < 0.05); and E, current disease state, in remission or not. Significance of treatment group clustering was determined by PERMANOVA (`adonis`) with 1,000 permutations. Bonferroni correction was applied for multiple comparisons.

Subjects in the MET group who were subsequently given AZ (MET/MET+AZ) after week 4, experienced a non-significant decrease in mean diversity at week 4 (week 0 vs week 4, Wilcoxon test, *P* = 0.07), consistent with this sub-group’s initial unresponsiveness to MET. However, after the addition of azithromycin at week 8, mean diversity decreased significantly (week 4 vs 8, Wilcoxon test, *P* ≤ 0.05), and became comparable to the level of the MET+AZ group at week 4 (Wilcoxon test, *P* = 0.2). Diversity remained low at week 12 in the MET/MET+AZ group, possibly because many subjects were still receiving their 8-week combination antibiotic regimen at that time.

Next, we examined the magnitude of the change in microbiota structure in response to antibiotics within each treatment group by calculating the median difference in unweighted UniFrac pairwise distance among samples from the same patient at different study timepoints. Microbiome composition in all three treatment groups changed to the same degree between weeks 0 and 4, reflected in a mean unweighted UniFrac pairwise distance of ~0.5, with no significant difference between groups (Fig. 2B). Thereafter, smaller changes occurred between weeks 4 and 8, but the MET/MET+AZ group showed a significantly larger shift in community structure than the MET group, possibly reflecting the addition of azithromycin (Fig. 2B).

We then assessed the contribution of current antibiotic use, treatment group, and remission status to variation in the data using principal coordinates analysis (PCoA) based on unweighted Unifrac distances.^25^ The use of antibiotics at the time of sample collection was a significant source of variation (PERMANOVA, corrected *P*-value < 0.01) (Fig. 2C), as was treatment group (PERMANOVA, corrected *P*-value < 0.05) (Fig. 2D) and remission status (PERMANOVA, corrected *P*-value < 0.05) (Fig. 2E). Other variables, such as gender, additional medications or dietary regimes had no significant impact on the microbiome composition (Table 2, Supplementary Fig. 3).

**Table 2.**
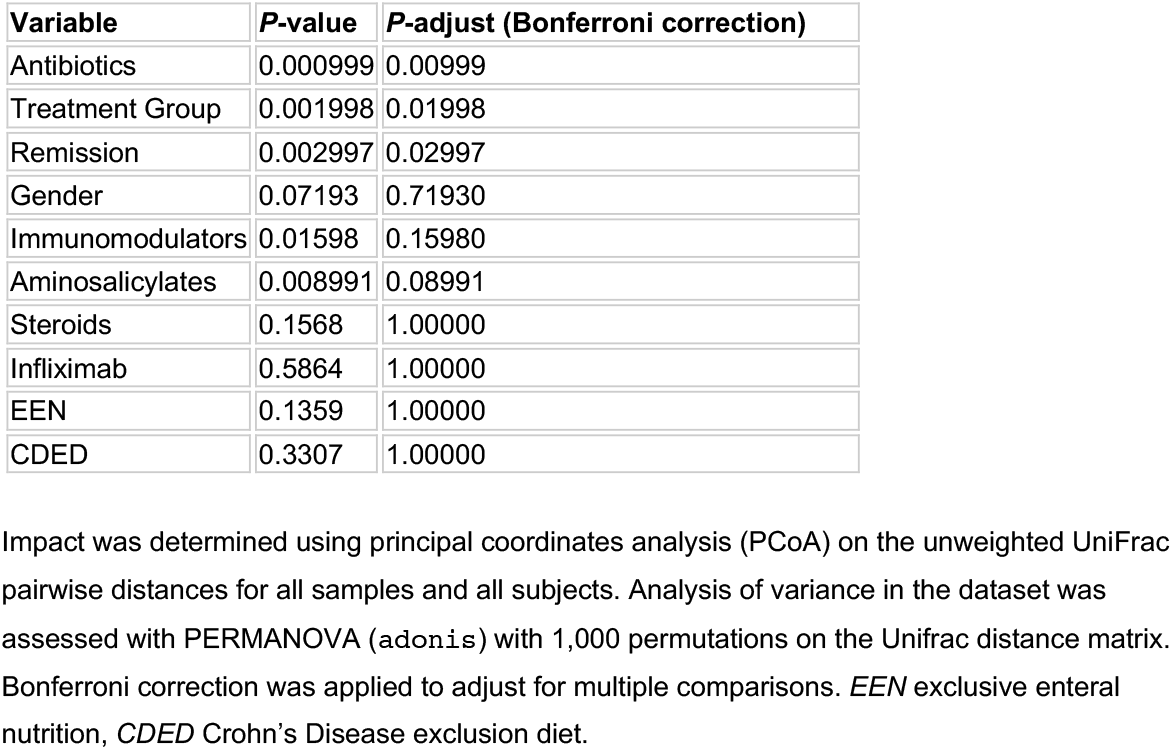
Impact of treatment, disease, and gender on microbiome composition.

| **Variable** | ***P*-value** | ***P*-adjust (Bonferroni correction)** |
| --- | --- | --- |
| Antibiotics | 0.000999 | 0.00999 |
| Treatment Group | 0.001998 | 0.01998 |
| Remission | 0.002997 | 0.02997 |
| Gender | 0.07193 | 0.71930 |
| Immunomodulators | 0.01598 | 0.15980 |
| Aminosalicylates | 0.008991 | 0.08991 |
| Steroids | 0.1568 | 1.00000 |
| Infliximab | 0.5864 | 1.00000 |
| EEN | 0.1359 | 1.00000 |
| CDED | 0.3307 | 1.00000 |

Impact was determined using principal coordinates analysis (PCoA) on the unweighted UniFrac pairwise distances for all samples and all subjects. Analysis of variance in the dataset was assessed with PERMANOVA (*adonis*) with 1,000 permutations on the Unifrac distance matrix. Bonferroni correction was applied to adjust for multiple comparisons. *EEN* exclusive enteral nutrition, *CDED* Crohn's Disease exclusion diet.

In order to identify groups of bacterial taxa with distinct responses to the different antibiotic regimens, we performed differential abundance testing between samples from weeks 0 and 4. ASVs with significant changes in abundance are summarized in a heat map, showing their relative abundance in each treatment group across all time points (Fig. 3). The top three clusters of the heatmap show ASVs that increased in abundance during the period of treatment and then decreased after antibiotics were withdrawn. While ASVs 7 and 61 (both *Enterococcus*) increased in abundance in both treatment groups, ASV 29 (*Streptococcus*) and ASV 27 (*Klebsiella*) increased in the MET group only. This can be explained by the known sensitivity of streptococci and *Klebsiella* to azithromycin and insensitivity to metronidazole.^26^

**FIGURE 3.**
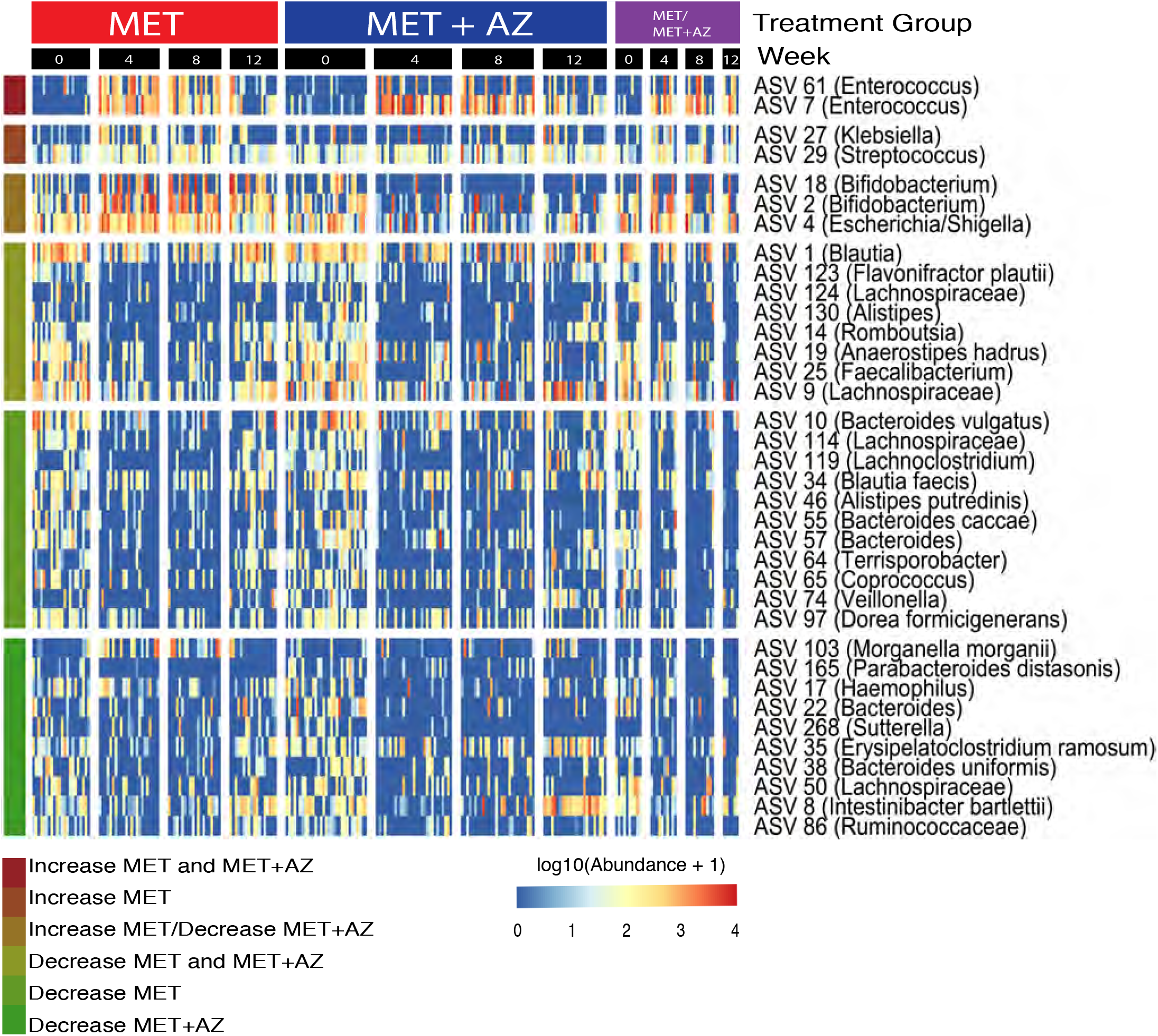
Microbial abundances shift in response to antibiotic exposure. ASVs were filtered for presence in at least 20% of samples. Testing for significant differences in ASV abundance between weeks 0 and 4 was performed using DeSeq2 in either the MET or MET+AZ treatment groups. Only ASVs with significant differences in abundance between weeks 0 to 4 are displayed. Abundances of ASVs were log_10_-transformed prior to plotting and partitioned by treatment group and week. Colored bars at the left end of the heatmap denote ASV responses to the treatments.

ASVs 18 and 2 (both *Bifidobacterium*) and ASV 4 (*Escherichia-Shigella*) increased in abundance in the MET group, but decreased in the MET+AZ group. The bottom three clusters of the heatmap include ASVs that decreased in abundance during single and/or combination antibiotic treatment and recovered at least partially after cessation of antibiotic administration (Fig. 3). ASVs 10 (*Bacteroides vulgatus*), 114 (*Lachnospiraceae*), 119 (*Lachnoclostridium*), 130 (*Alistipes*), 3 (*Faecalibacterium prausnitzii*), 34 (*Blautia faecis*), 46 (*Alistipes putredinis*), 55 (*Bacteroides caccae*), 57 (*Bacteroides*), 64 (*Terrisporobacter*), 65 (*Coprococcus*), 74 (*Veillonella*) and 97 (*Dorea formicigenerans*) significantly decreased in abundance in the MET treatment group, while ASVs 103 (*Morganella morganii*), 165 (*Parabacteroides distasonis*), 17 (*Haemophilus*), 22 (*Bacteroides*), 25 (*Faecalibacterium*), 268 (*Sutterella*), 35 (*Erysipelatoclostridium ramosum*), 38 (*Bacteroides uniformis*), 50 (*Lachnospiraceae*) and 86 (*Ruminococcaceae*) decreased significantly in the MET+AZ group.

### Microbiota and Crohn’s Disease remission

To understand better how antibiotics might influence microbiota structure and remission, we calculated pairwise Bray-Curtis dissimilarity scores for remission samples and compared those to the pair-wise scores for non-remission samples in each treatment group. Week 4 and 8 samples from patients in the MET group who were in remission at that time were significantly more similar to each other than they were to week 4 and 8 samples from patients who were not in remission at the time (Wilcoxon text, *P* ≤ 0.001) (Fig. 4A). We observed no significant differences for samples from the MET+AZ group. Furthermore, we found that remission samples from weeks 4 and 8 formed distinct clusters reflecting treatment group (PERMANOVA, *P* ≤ 0.001), a pattern that was not observed for non-remission samples (Fig. 4B).

**FIGURE 4.**
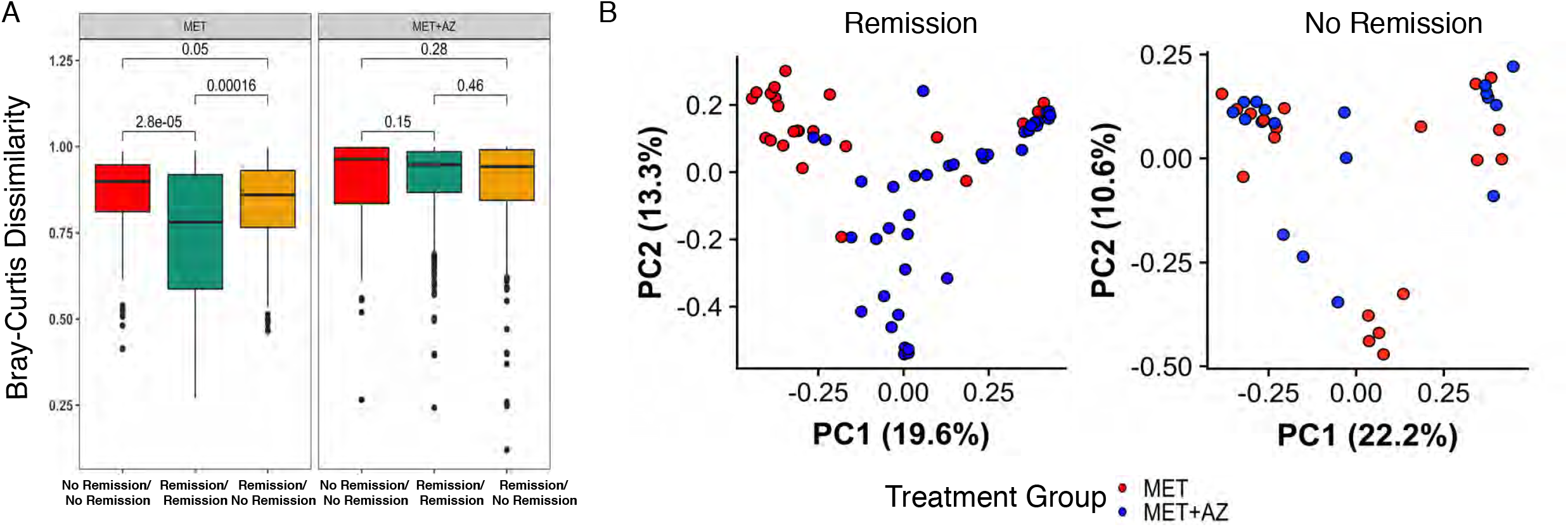
Microbiota at time of remission reflects antibiotic exposure. **A,** Beta-diversity measured by Bray-Curtis dissimilarity between samples in remission and non-remission among MET and MET+AZ subjects during antibiotic treatment (weeks 4 and 8). **B,** PCoA on the Bray-Curtis dissimilarity of stool samples collected during antibiotic treatment (weeks 4 and 8). Remission samples (PERMANOVA, *P* = 0.0009), no remission samples (PERMANOVA, *P* = 0.1848). Significance of treatment group clustering was determined by PERMANOVA (`adonis`) with 1,000 permutations.

Based on these observations, we sought to identify treatment-specific microbial indicators of CD remission. We used Random Forest modeling to predict whether a patient was in remission based on their microbiota structure at that time. First, we constructed a model based on the abundances of ASVs found in remission and non-remission samples collected during antibiotic treatment (Model 1, weeks 4 and 8). Because the numbers of remission and non-remission samples in the MET+AZ group were too uneven to be able to construct a useful model, we focused on the MET group only. This model classified remission in the MET patients with an AUC of 0.777 (95% CI, 0.3229, 0.8366; sensitivity, 0.8750; specificity, 0.2857; *P* = 0.4006), and classified remission in the MET+AZ group with an AUC of 0.68 (95% CI, 0.4066, 0.6764; sensitivity, 0.3846; specificity, 0.8889; *P* = 0.9909) (Fig. 5A), but did not achieve statistical significance in either case. In order to test an alternative model that incorporated a greater number of non-remission samples, we constructed Model 2 with samples from week 0 as well; these samples represent an alternative non-remission state prior to antibiotic administration, but at the same time, these samples introduce antibiotic treatment as a confounding factor (Model 2, weeks 0, 4 and 8). Model 2 classified remission in MET patients with improved accuracy and an AUC of 0.879 (95% CI: 0.683, 0.9877; sensitivity: 0.7778; specificity: 1.000; *P* < 0.001) and remission in the MET+AZ group with a lower AUC of 0.695 (95% CI: 0.5038, 0.7156; sensitivity: 0.20513; specificity: 0.93878; *P* = 0.1672) (Fig. 5A).

**FIGURE 5.**
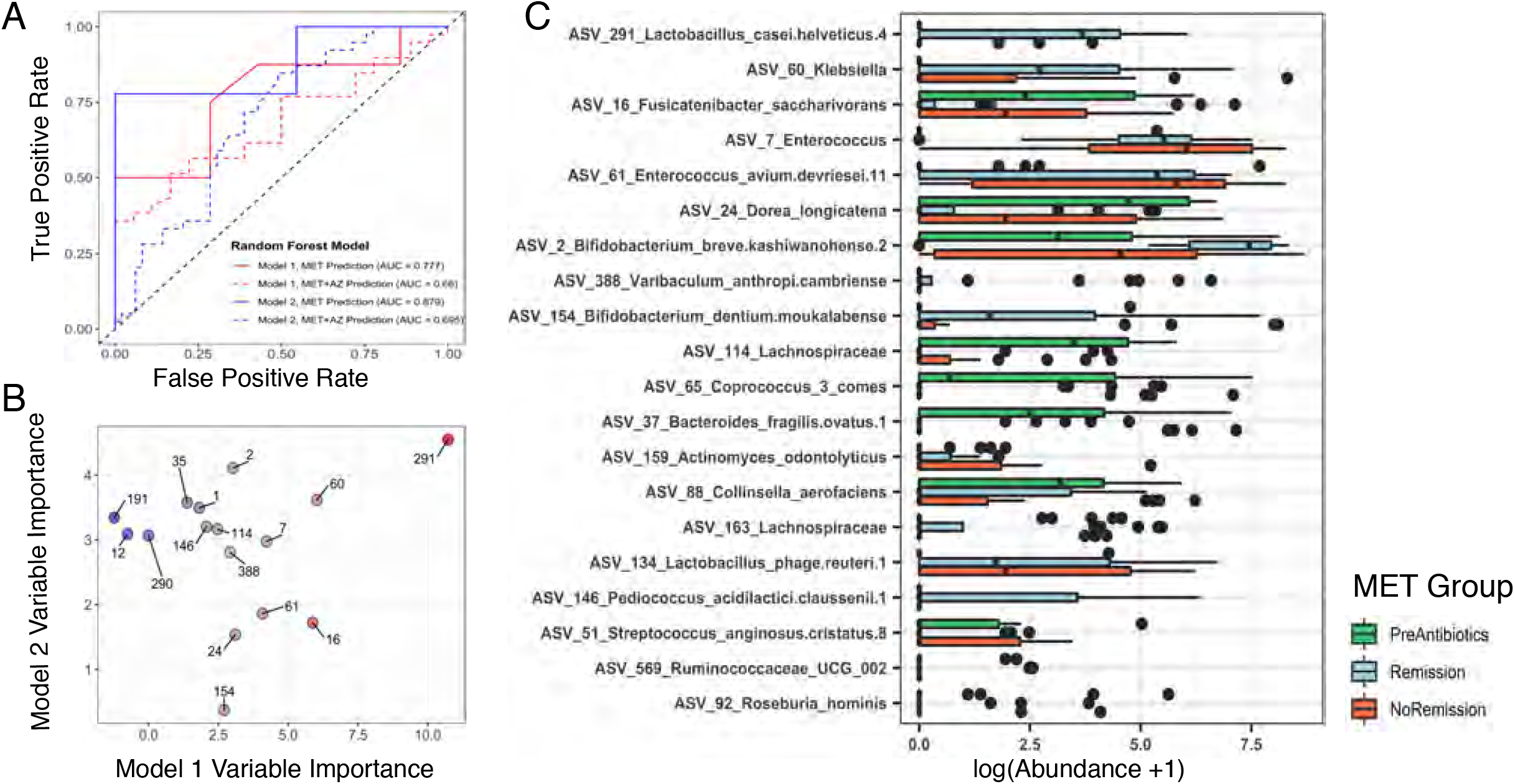
Random Forest models classify disease remission from microbiota profiles. **A,** The ROC curves indicate the accuracy of the random forest classification models built using microbiota data. Color of lines indicates the dataset on which the model was trained (Model 1, MET weeks 4 and 8, red; Model 2, MET weeks 0, 4 and 8, blue). Solid lines indicate accuracy of the model in classifying remission in patients from the same treatment group (MET), while the dashed lines indicate the accuracy of each model for classifying remission in patients from the MET+AZ treatment group. *AUC* Area under the curve. **B,** Variable importance values for the 10 most important ASVs used to build each random forest model are plotted against each other. Points are labeled by ASV number. Point color indicates the importance score (Model 1 specific, red; Model 2 specific, blue; both models, gray). **C,** Log-transformed abundances of top 20 ASVs that were of common importance for both Random Forest models.

Models 1 and 2 each assigned importance scores to each ASV based on the increase in model prediction error when that feature was randomly permuted while all others were left unchanged. The 10 most important ASVs in each model are displayed in Figure 5B. The most important ASV for both models was ASV 291 (*Lactobacillus*) (Fig 5B). The ASVs with high importance scores had different abundances in pre-antibiotics, remission, and non-remission samples. Compared to their relative abundances in pre-antibiotic samples, we observed a large increase in ASV 291 (*Lactobacillus*), ASV 60 (*Klebsiella*) and ASVs 2 and 154 (both *Bifidobacterium*), specifically in remission samples (Fig. 5C). In agreement with the results shown in Figure 3, we observed an increase in abundance of ASVs 7 and 61 (both *Enterococcus*) from the pre-antibiotic state, but no difference between remission and nonremission samples. ASVs that decreased in remission samples, but not in non-remission samples, included ASV 16 (*Fusicatenibacter saccharivorans*) and ASV 24 (*Dorea longicatena*).

We wondered whether there were microbial signatures present prior to antibiotic administration that would predict remission after treatment. Although not statistically significant, pre-antibiotic samples from MET subjects who were in remission at week 4 trended towards higher alpha diversity (see Supplementary Fig. 4A, as Supplementary Data Content 4), a pattern reported in other prediction-focused studies.^27,28^ Due to the limited number of preantibiotic samples available, we combined samples from both treatment groups, and information on each subject’s age, sex, disease duration, tissue involvement, and the current disease state. Unfortunately, we still were not able to construct a robust model (AUC = 0.8; 95% CI: 0.4099, 0.8666; sensitivity, 0.8; specificity, 0.5; *P* = 0.24) (see Supplementary Fig. 4B, as Supplementary Data Content 4). However, we identified ASV clusters with higher mean abundance in pre-antibiotic samples of subjects that went on to achieve remission, such as Cluster 8 (*Peptostreptococcaceae*), Cluster 24 (*Alistipes*), Cluster 23 (*Clostridium* sensu stricto) and Cluster 27 (*Parabacteroides*) (see Supplementary Fig. 4C, as Supplementary Data Content 4). In contrast, for example, ASV Cluster 61 (*Fusobacterium*) was more abundant in preantibiotic samples of subjects who did not subsequently achieve remission, in accordance with other studies.^7–9^

## DISCUSSION

In this study, we determined the effects of MET-only and combination MET+AZ therapy on the gut microbiota in pediatric CD patients. We demonstrated a treatment-specific effect of both antibiotic regimens on microbiota structure, especially at the time of clinical disease remission. We assessed the utility of microbiota signatures for classifying disease remission as well as the capacity of baseline (pre-antibiotic) microbiota structure to predict future treatment response.

In comparison to previous work, the strength of our study lies in the use of samples from a multinational cohort and from CD patients exclusively. Small cohort size is a common problem and has led to co-evaluation of samples from mixed patient populations with CD and ulcerative colitis, while recent data highlight the differences between those subtypes of IBD with respect to the microbiota.^29^ Second, many previous studies have characterized the microbiota of patients based on one sample and time point,^7,9^ while the design of this study provided an opportunity to monitor the microbiota before, during, and after treatment, which we believe is critical for identifying microbial signatures related to different clinical outcomes.

We acknowledge weaknesses in study design, such as failed randomization of gender and subjects with L1 Paris classification, as well as observed impact of 5ASA on baseline microbiome, although distribution of 5ASA between treatment groups was equal. The use of additional medication by some patients during the study was also a potential confounding factor. The switching between treatment arms upon failure to achieve remission also complicated our analysis. These factors are difficult to navigate in a patient population with such a complex disease and individual treatment requirements. Furthermore, our ability to build a robust predictive model for both treatment arms was limited by the sample size.

It is well established that antibiotic treatment creates an ecosystem-wide disturbance and decreases the overall diversity of the gut microbiome.^30^ To our knowledge, few studies have examined the effects of single versus combination antibiotics on the gut microbiome.^31^ While we observed a general decrease in diversity in response to antibiotics, our results also showed a distinct impact on microbiota structure and a shift in abundance of specific bacterial ASVs in each treatment group. These findings highlight the need to understand better the additive, antagonistic, synergistic, and non-linear effects of antibiotic treatments with regards to patient outcome. Despite different effects on the gut microbiota, the two treatments both produced remission-compatible microbiota configurations.

Metronidazole is one of the most prescribed antibiotics in the treatment of pediatric CD, but to our knowledge, there has been no in-depth study of the effects of metronidazole treatment on the gut microbiota in humans to date. One study in mice, comparing the impact of metronidazole versus streptomycin on the microbiota and subsequent *C. rodentium-induced* colitis, observed treatment-specific effects, not only on microbiota structure, but on gut mucosal immune responses as well.^32^ In the absence of *C. rodentium* infection, metronidazole was found to decrease goblet cell expression of MUC2, thereby leading to a thinning of the inner mucus layer, which could be detrimental for intestinal homeostasis and promote chronic intestinal inflammation. Although this observation has not been confirmed in humans, it has important implications for the use of MET in IBD patients.

Recent studies reported superior outcomes for MET plus AZ, as compared to MET alone.^13,14^ AZ penetrates multiple intestinal compartments^15,16^ and enables targeting of adherent and invasive *E. coli* (AIEC) strains and other bacteria that have been associated with CD at disease onset in children.^9,17–19^ Consistent with previous reports, we observed reduced abundances of ASV 4 (*Escherichia/Shigella*) in the MET+AZ group, but increased abundances in the MET group, which may reflect activity of azithromycin against this taxon in our cohort. To assess the presence of AIEC specifically, patient biopsies could be used in the future to access the mucosa-adherent or intracellular microbial constituents.

We also observed that some bacterial taxa such as *Enterococcus, Streptococcus, Klebsiella, Bifidobacterium* and *Enterobacteriaceae* increased in abundance during antibiotic treatment. This could be due to creation or expansion of nutritional and/or environmental niches during or after general community disturbance or could indicate intrinsic or acquired antibiotic resistance. Our finding of increased abundances of *Enterococcus* ASVs in both treatment groups is concerning, as they possess multiple mechanisms for resisting antibiotics; their expansion could predispose patients to invasive infections.^33,34^

An important question is whether the microbiome itself might be used as a therapeutic agent for Crohn’s disease, for example by administration of specific beneficial microbes as probiotics.^35^ Our study identified multiple *Lactobacillus* ASVs that were important for the classification of clinical remission and whose abundance significantly increased during remission. *Lactobacillus* species are commonly used as probiotics and several *in vitro* and animal studies suggest that they may reduce inflammation in CD.^36,37^ In humans, clinical trials with *Lactobacillus* probiotics in CD patients have so far been unsuccessful,^38,39^ highlighting the need to understand not only the biology of *Lactobacillus* species but microbial community ecology.

Two recent studies used microbiota data to predict the success of biologics in adult IBD patients and identified taxa associated with disease remission. Ananthakrishnan *et al.* reported a decrease in abundance of five taxa associated with remission in response to anti-integrin biologic therapy - among them *Bifidobacterium longum*.^40^ In contrast, we observed a large increase in abundance of two *Bifidobacterium* ASVs in antibiotic-associated remission samples. Doherty *et al.* showed that the relative abundance of *Escherichia/Shigella* was lower in subjects in remission associated with ustekinumab therapy, than it was in subjects with active CD.^27^ In our study subjects, we observed decreased abundance of an *Escherichia/Shigella* ASV in the MET+AZ group, which achieved higher remission rates, although the ASV was not implicated in our predictive model of remission. Both studies reported higher microbiota alpha diversity at baseline in patients who later achieved remission. Although not statistically significant, baseline samples from MET subjects in our cohort who achieved remission at week 4 trended towards higher alpha diversity as well.

We suggest that Random Forest models can be useful for microbiota-based classification and prediction. Our efforts in constructing useful classifiers for disease remission or prediction of treatment response to antibiotics draws attention to the importance and difficulties of obtaining sufficient numbers of subjects and samples. Moreover, the identification of remission-associated patterns may depend on the cohort characteristics, such as age and disease duration, as well as on the therapy, its dosage and duration.

## CONCLUSION

There are several important implications of our findings. Our results suggest that there is no single antibiotic-associated remission state, but that different regimens may impact the microbiome in a distinct manner, yet lead to remission. We show that the classification of pediatric CD patients in antibiotic-associated clinical remission based on their microbiota structure is possible, although large sample sizes will be required for accurate model construction. We suggest that classification and prediction models based on microbiota signatures may be another tool for monitoring disease and treatment response. Moreover, such models may also assist in the development and testing of more precise approaches for microbiome manipulation and might eventually lead to more effective management of CD and other forms of inflammatory bowel disease.

## ACKNOWLEDGEMENTS

We are grateful to the study participants. We thank Alvaro Hernandez at the University of Illinois Roy J. Carver Biotechnology Center for outstanding DNA sequencing services. This research was supported by National Science Foundation Graduate Research Fellowship DGE-114747 (D.S.), National Institute of General Medical Sciences of the National Institutes of Health training grant T32GM007276 (D.S.), Helmsley Foundation grant 2014PG-IBD014 (D.A.R.), Thomas C. and Joan M. Merigan Endowment at Stanford University (D.A.R.), and Chan Zuckerburg Biohub Microbiome Initiative (D.A.R.).

---

**Supplementary Figure 1.**
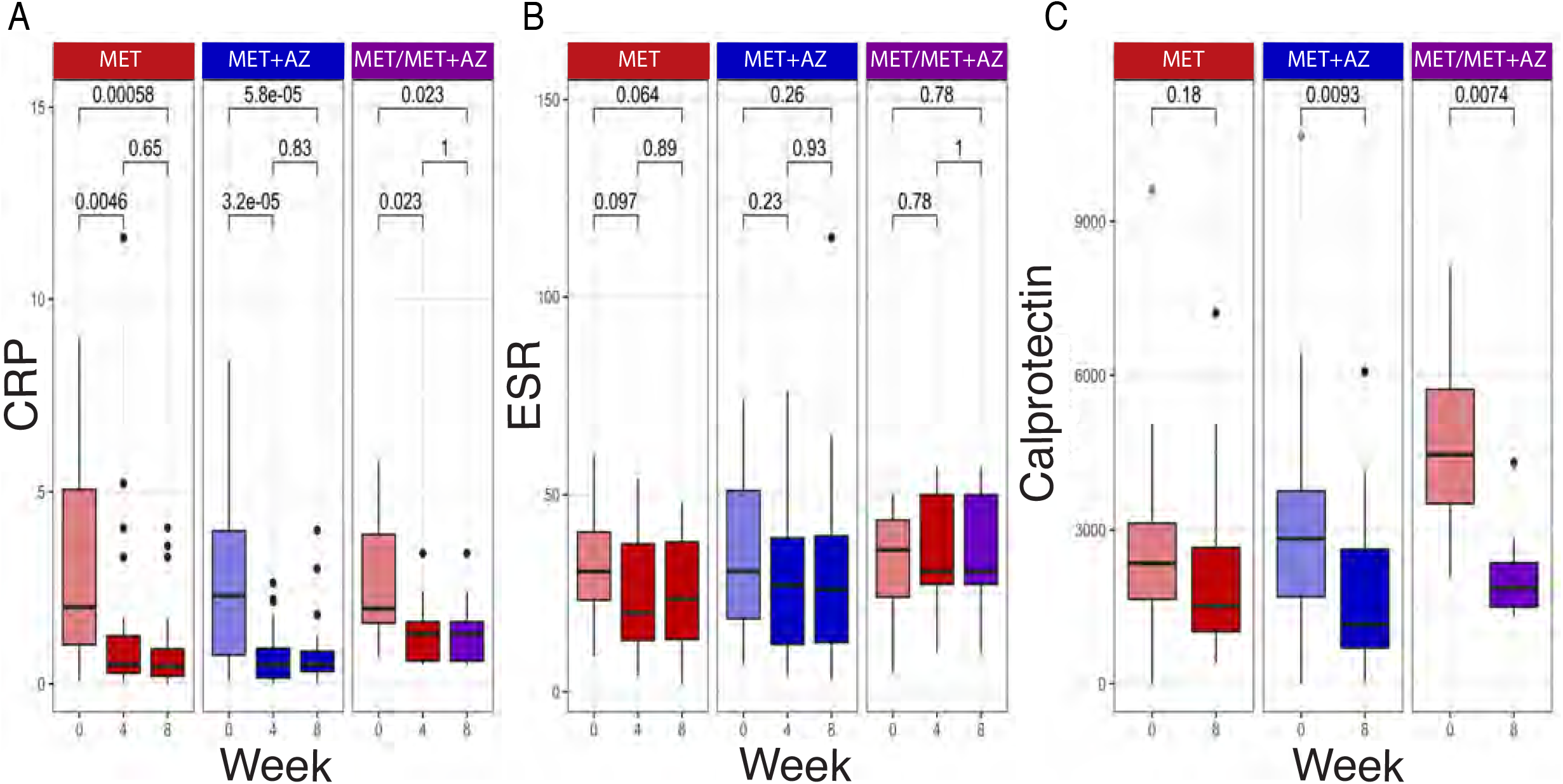
Change in markers of inflammation following antibiotic treatment. **A,** C-reactive protein (CRP) levels at weeks 0, 4, and 8, colored by treatment group (MET, red; MET+AZ, blue; MET/MET+AZ, purple). **B,** Fecal calprotectin levels at weeks 0, and 8. **C,** Erythrocyte sedimentation rate (ESR) at weeks 0, 4, and 8. Samples collected during antibiotic administration are darker in color. *P*-values between time points within a treatment group are displayed (Wilcoxon Test).

**Supplementary Figure 2.**
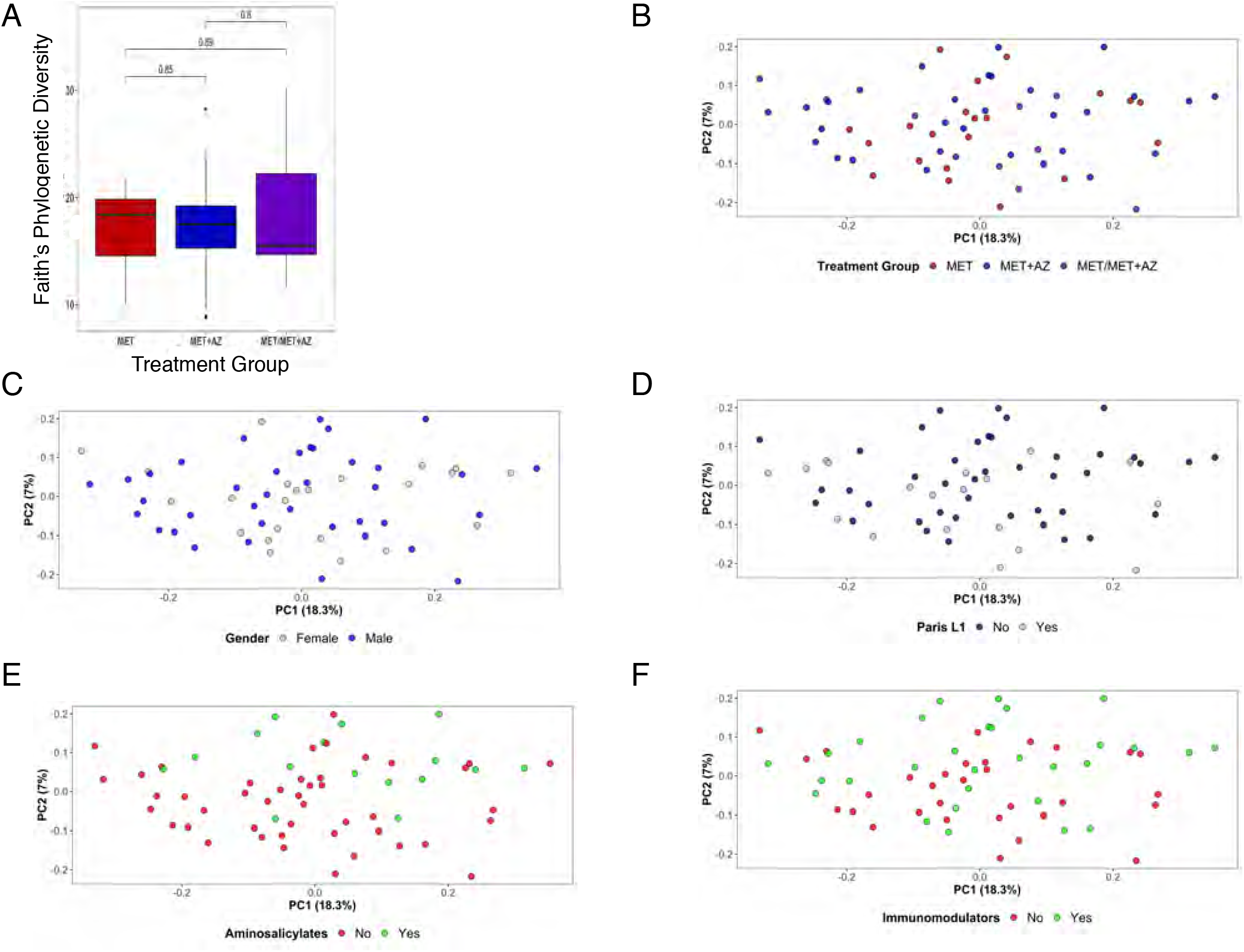
Baseline microbiota diversity and composition. A, Faith’s phylogenetic alpha diversity of samples collected at baseline (week 0) was compared between treatment groups (Wilcoxon test). B-F, PCA was performed on the unweighted UniFrac pairwise distances among all stool samples at baseline (B, Treatment Group, *P* = 1; C, Gender, *P* = 1; D, L1 Paris Classification, *P* = 0.34; E, 5-aminosalicylic acid, *P* < 0.05; F, immunomodulators, *P* = 0.11). Significance of group clustering was determined by PERMANOVA (`adonis`) with 1,000 permutations. Bonferroni correction was applied for multiple comparisons.

**Supplementary Figure 3.**
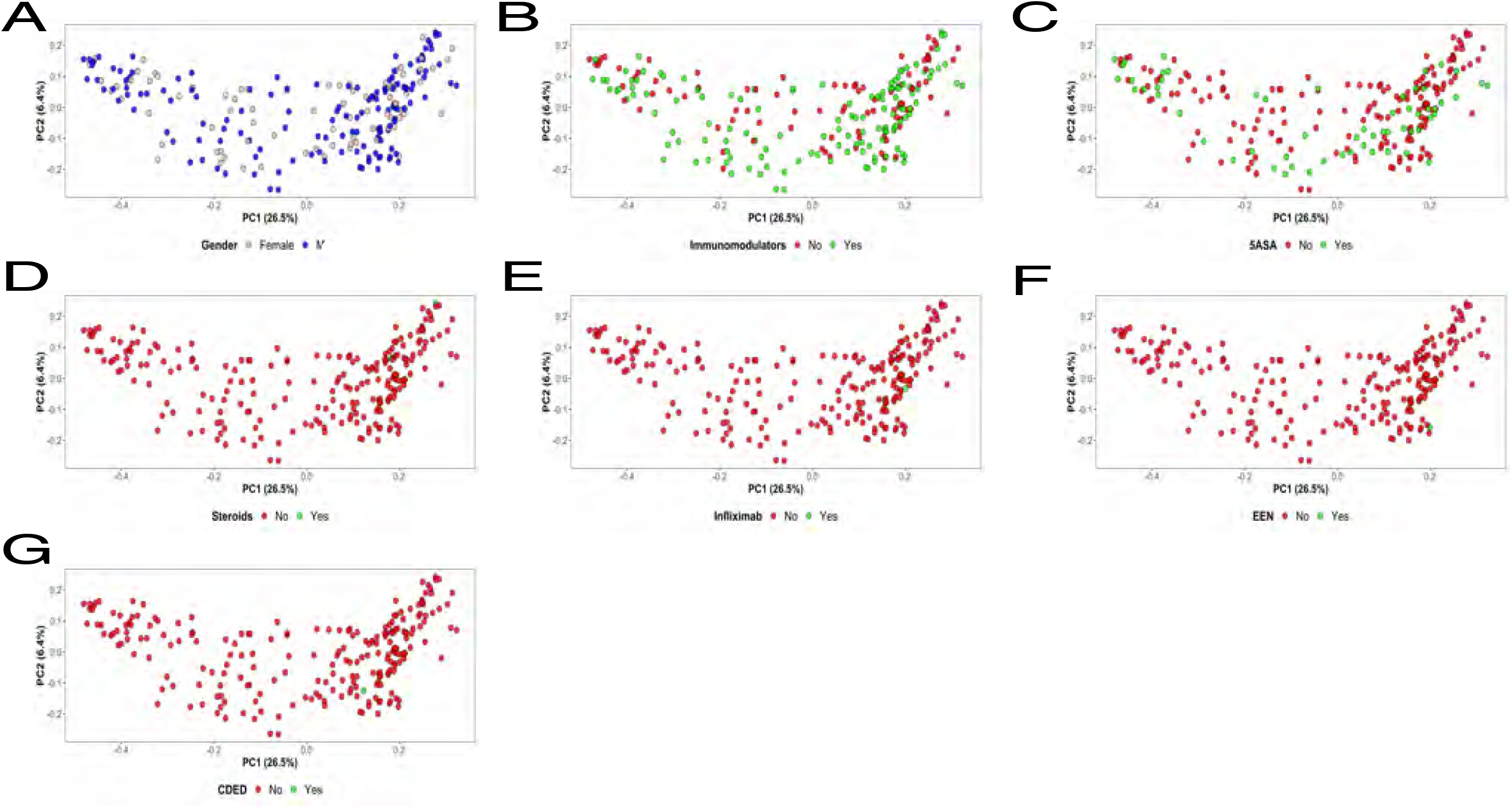
Impact of gender, additional medications and diet regimes on microbiome composition. Principal coordinates analysis (PCoA) on the unweighted UniFrac pairwise distances among all samples and subjects. Samples are colored by A, gender (PERMANOVA, adjusted *P* = 0.72); B, immunomodulators (PERMANOVA, adjusted *P* = 0.16); C, aminosalicylates (PERMANOVA, adjusted *P* = 0.09); D, steroids (PERMANOVA, adjusted *P* = 1); E, infliximab (PERMANOVA, adjusted *P* = 1); F, exclusive enteral nutrition (EEN) (PERMANOVA, adjusted *P* = 1); G, Crohn’s disease exclusion diet (CDED) (PERMANOVA, adjusted *P* = 1). Significance of treatment group clustering was determined by PERMANOVA (`adonis`) with 1,000 permutations. Bonferroni correction was applied for multiple comparisons.

**Supplementary Figure 4.**
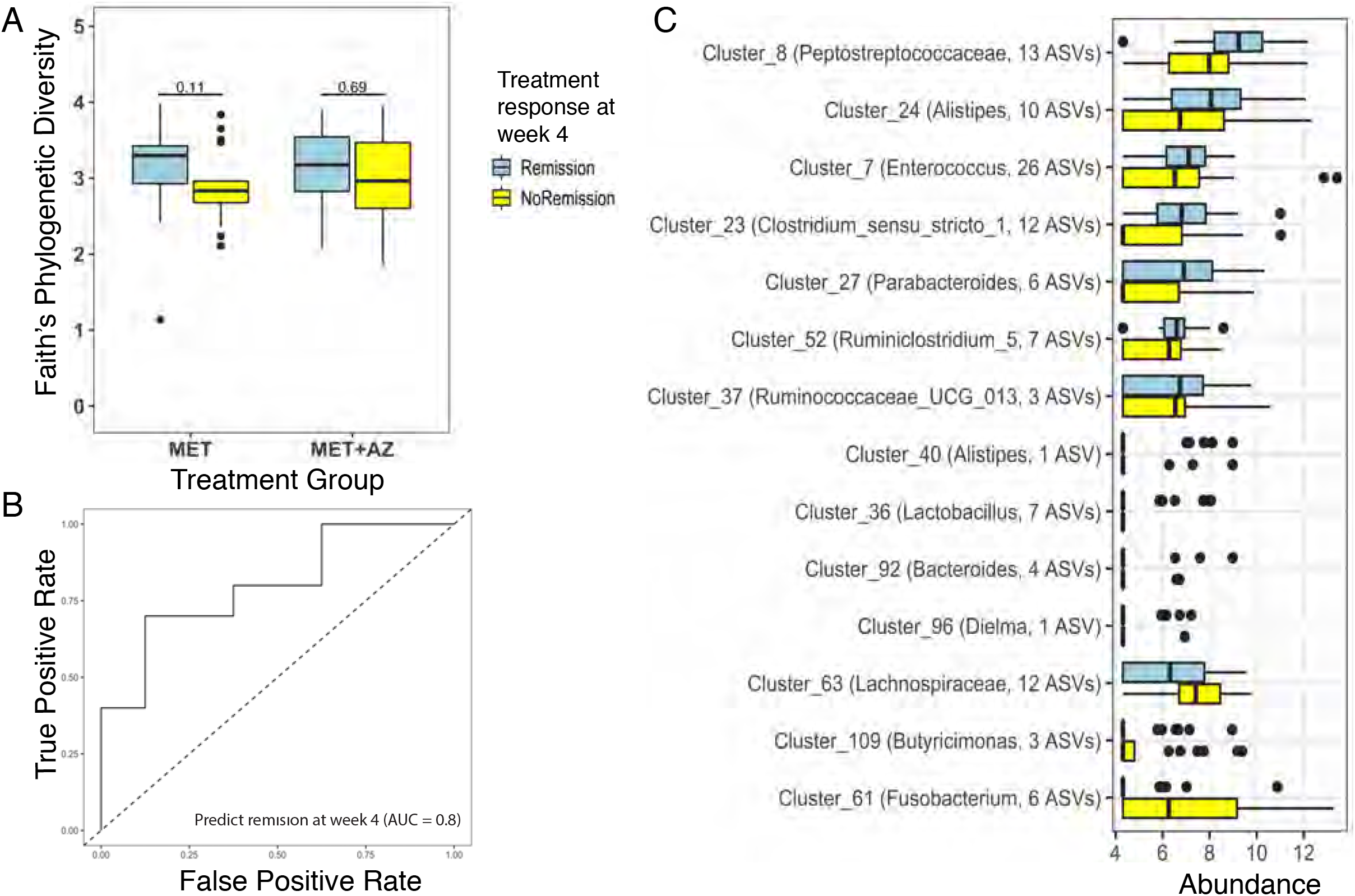
Pre-treatment microbiota structure predicts treatment outcome. **A,** Alpha-diversity of samples collected at baseline (week 0) grouped by their treatment group and remission status at week 4 (Wilcoxon Test). **B,** The gray ROC curve indicates the accuracy of the random forest classification model built using microbiome data from week 0 as well as gender, age, disease duration, Paris classification, pre-antibiotic immunomodulators, PCDAI and CRP to predict response to treatment at week 4. *AUC* Area under the curve. **C,** Abundances of ASV clusters that are important for the remission-forecasting random forest models. Abundances were transformed using a variance stabilizing transformation (Bioconductor package `vsn`).

## Notes

This research was funded by National Science Foundation Graduate Research Fellowship DGE-114747 (D.S.), National Institute of General Medical Sciences of the National Institutes of Health training grant T32GM007276 (D.S.), Helmsley Foundation grant 2014PG-IBD014 (D.A.R.), Thomas C. and Joan M. Merigan Endowment at Stanford University (D.A.R.), and Chan Zuckerburg Biohub Microbiome Initiative (D.A.R.)

#### Summary of Updates

Figure 1 has been revised, Table 1 has been revised, Table 2 is new, Supplementary Figure 2 is new.

